# *AtRsgA* from *Arabidopsis thaliana* controls maturation of the small subunit of the chloroplast ribosome

**DOI:** 10.1101/225052

**Authors:** Marcin Janowski, Reimo Zoschke, Lars Scharff, Silvia Martinez Jaime, Camilla Ferrari, Sebastian Proost, Nooshin Omranian, Magdalena Musialak-Lange, Zoran Nikoloski, Alexander Graf, Mark A. Schöttler, Arun Sampathkumar, Neha Vaid, Marek Mutwil

**Affiliations:** Max Planck Institute of Molecular Plant Physiology, Am Muehlenberg 1, 14476 Potsdam, Germany; Copenhagen Plant Science Centre, Department of Plant and Environmental Sciences, University of Copenhagen, Thorvaldsensvej 40, DK-1871 Frederiksberg C, Denmark; Bioinformatics Group, Institute of Biochemistry and Biology, University of Potsdam, Karl-Liebknecht-Str. 24-25, 14476 Potsdam-Golm, Germany; School of Biological Sciences, Nanyang Technological University, 60 Nanyang Drive, Singapore 637551, Singapore

## Abstract

**Summary:** Plastid ribosomes are very similar in structure and function to ribosomes of their bacterial ancestors. Since ribosome biogenesis is not thermodynamically favourable at biological conditions, it requires activity of many assembly factors. Here, we have characterized a homolog of bacterial *rsgA* in *Arabidopsis thaliana* and show that it can complement the bacterial homolog. Functional characterization of a strong mutant in Arabidopsis revealed that the protein is essential for plant viability, while a weak mutant produced dwarf, chlorotic plants that incorporated immature pre-16S ribosomal RNA into translating ribosomes. Physiological analysis of the mutant plants revealed smaller, but more numerous chloroplasts in the mesophyll cells, reduction of chlorophyll *a* and *b*, depletion of proplastids from the rib meristem and decreased photosynthetic electron transport rate and efficiency. Comparative RNA-sequencing and proteomic analysis of the weak mutant and wild-type plants revealed that various biotic stress-related, transcriptional regulation and post-transcriptional modification pathways were repressed in the mutant. Intriguingly, while nuclear- and chloroplast-encoded photosynthesis-related proteins were less abundant in the mutant, the corresponding transcripts were upregulated, suggesting an elaborate compensatory mechanism, potentially via differentially active retrograde signalling pathways. To conclude, this study reveals a new chloroplast ribosome assembly factor and outlines the transcriptomic and proteomic responses of the compensatory mechanism activated during decreased chloroplast function.

**Significance statement:** AtRsgA is an assembly factor necessary for maturation of the small subunit of the chloroplast ribosome. Depletion of AtRsgA leads to dwarfed, chlorotic plants and smaller, but more numerous chloroplasts. Large-scale transcriptomic and proteomic analysis revealed that chloroplast-encoded and - targeted proteins were less abundant, while the corresponding transcripts were upregulated in the mutant. We analyse the transcriptional responses of several retrograde signalling pathways to suggest a mechanism underlying this compensatory response.

## Introduction

Plastids have retained many prokaryotic features from its bacterial ancestor, such as the presence of operons and polycistronic RNAs for most of the chloroplast-encoded genes. These genes are transcribed by a bacteria-type, chloroplast-encoded RNA polymerase and nucleus-encoded sigma factors of the s70 type that recognize promoters resembling bacterial promoters (Liere and Börner, 2007). Chloroplast ribosomes share many features with bacterial ribosomes as ribosomal RNAs (rRNAs) and 21 ribosomal proteins from the small subunit (30S) have clear orthologs in *Escherichia coli*. Similarly, the large ribosomal subunit (50S) composed of three rRNAs (4.5S rRNA, 5S rRNA and 23S rRNA) and 31 ribosomal proteins have orthologs in *E.coli*. Interestingly, 12 out of 21 proteins associated with 30S and 22 out of 31 proteins associated with 50S ribosomal subunits are encoded by the nucleus rather than the chloroplast genome (Yamaguchi and Subramanian, 2000). Many of these proteins are essential in *E.coli* and plants (Rogalski *et al*., 2006; Fleischmann *et al*., 2011; Tiller and Bock, 2014). Also, several ribosome assembly factors, tRNAs, initiation, elongation, and termination factors are conserved between plastids and *E.coli* (Britton, 2009; Alkatib *et al*., 2012; Meurer *et al*., 2002; Albrecht *et al*., 2006; Motohashi *et al*., 2007; Shen *et al*., 2013).

Ribosomal biogenesis is among the most energy consuming processes of the cell, and involves stepwise processing of the primary rRNA transcript and the correct assembly of ribosomal proteins and rRNAs into functional ribosomal subunits. This highly coordinated process is facilitated by many trans-acting factors, including accessory proteins and small RNAs (Fromont-Racine *et al*., 2003; Kaczanowska and Rydén-Aulin, 2007). *In-vitro* reconstitution of ribosomes without trans-acting factors requires high ionic strength, heating and prolonged incubation time (Nierhaus and Dohme, 1974; Traub and Nomura, 1969), indicating the essential role of these factors under physiological conditions. Not surprisingly, more than half of these trans-acting factors are ATP- and GTP-binding proteins (rATPases and rGTPases) that provide energy for processes such as breaks and inter/intramolecular rearrangements, thus allowing association and dissociation of other assembly factors or ribosomal proteins for sequential progress of the ribosome assembly (Shajani *et al*., 2011).

Since many plastid ribosomal proteins and assembly factors are conserved between chloroplast and *E.coli*, and many of these proteins can be integrated into *E.coli* ribosomes, a great deal of information regarding bacterial ribosome biogenesis can be conveyed to the chloroplast ribosomes (Bubunenko and Subramanian, 1994; Weglohner *et al*., 1997). The detailed assembly maps for both ribosomal subunits and the necessary assembly factors needed for rRNA processing and/or ribosome maturation from *E.coli* are conserved in chloroplasts, despite more than 1.6 billion years since the endosymbiosis of the cyanobacterial ancestor of chloroplast (Dabbs, 1991; Talkington *et al*., 2005; Kaczanowska and Rydén-Aulin, 2007; Woodson, 2008; Connolly and Culver, 2013; Fournier *et al*., 2010; Davies *et al*., 2010; Jacob *et al*., 2013). However, since more than 90 *%* of endosymbiont bacterial genes migrated to host cell nuclear genome (Sato, 1999; Timmis *et al*., 2004), all of the ribosome biogenesis factors are nuclear-encoded. Consequently, their site of action in a plant cell (chloroplast or mitochondrion or both) remains to be determined.

RsgA is among the most extensively studied *E.coli* rGTPases. The null mutants of RsgA in *E.coli* and *Bacillus subtillis* show decreased growth rates, decreased levels of mature 70S ribosomes, and accumulation of 17S RNA (precursor of 16S RNA), indicating its role in the biogenesis of the 30S ribosomal subunit (Campbell *et al*., 2005; Himeno *et al*., 2004). However, its *Arabidopsis thaliana* homolog null mutant shows embryo lethality (Tzafrir *et al*., 2004), indicating that the assembly pathways of 70S ribosomes in bacteria and chloroplasts might have some subtle differences. In general, the rGTPases associate with the immature ribosomal subunits in GTP-bound form, resulting in enhanced GTPase activity, followed by dissociation of GDP-bound rGTPase from the subunit (Goto *et al*., 2011). Structurally, most rGTPases possess a nucleotide binding site and a GTPase domain. The latter consists of five conserved motifs; G1 to G5 (Leipe *et al*., 2002). The GTPase domain of several rGTPases is organized in a non-canonical order of G4-G5-G1-G2-G3, often referred to as circular permutated GTPase domain (Leipe *et al*., 2002; Elias and Novotny, 2008). Structural studies on immature 30S subunit from *ΔrsgA E.coli* revealed severe distortion at the decoding center of the 30S subunit, which makes it unable to associate with the 50S subunit to carry out mRNA translation (Jomaa *et al*., 2011). A recent study reports that *ΔrsgA* in *E.coli*, the immature 30S subunit can still assemble with a mature 50S subunit, although at a slower rate, explaining the slower growth rate phenotype (Thurlow *et al*., 2016). However, it’s function and site of action in plants remains unclear, which prompted us to study Arabidopsis thaliana RsgA (AtRsgA) gene in detail.

## Results

### AtRsgA can complement the small ribosomal subunit maturation deficiency in an *E. coli ΔrsgA* mutant

*E.coli* RsgA is involved in biogenesis of the small ribosomal subunit (30S). Its null mutant is characterized by slow growth under favourable growth conditions (Kimura *et al*., 2008; Goto *et al*., 2011). We identified homologs of *E.coli* RsgA in a cyanobacterium, a green alga, and a plant, demonstrating that RsgA belongs to a conserved protein family (Figure S1). The multiple protein sequence alignment revealed a conserved oligonucleotide/oligosaccharide binding motif (OB) and a GTPase domain. We also observed a plant-specific sequence insertion within the GTPase domain (Figure S1, marked in red box).

To investigate if the *Arabidopsis thaliana* RsgA homolog (AtRsgA) protein is also involved in 30S maturation, we transformed *E.coli ΔrsgA* null mutant with full-length *AtRsgA* cDNA. We observed that *AtRsgA* can partially rescue the growth phenotype in liquid (Figure 1A) and solid media (Figure 1B), when compared to *ΔrsgA* and *ΔrsgA* transformed with empty vector (EV). To confirm if the rescue of slow growth phenotype is due to improved 30S subunit biogenesis, we quantified ribosomal RNAs (rRNAs) which are indicative of the amount of their respective ribosomal subunits (Tiller *et al*., 2012)(Figure S2). We observed an increased ratio of 16S rRNA (30S subunit) to 23S rRNA (50S subunit) in *AtRsgA* complemented *ΔrsgA* as compared to the controls (Figure 1C). This demonstrated that AtRsgA can partially rescue the 16S maturation deficiency and confirmed that AtRsgA is a 30S assembly factor.

**Figure 1.**
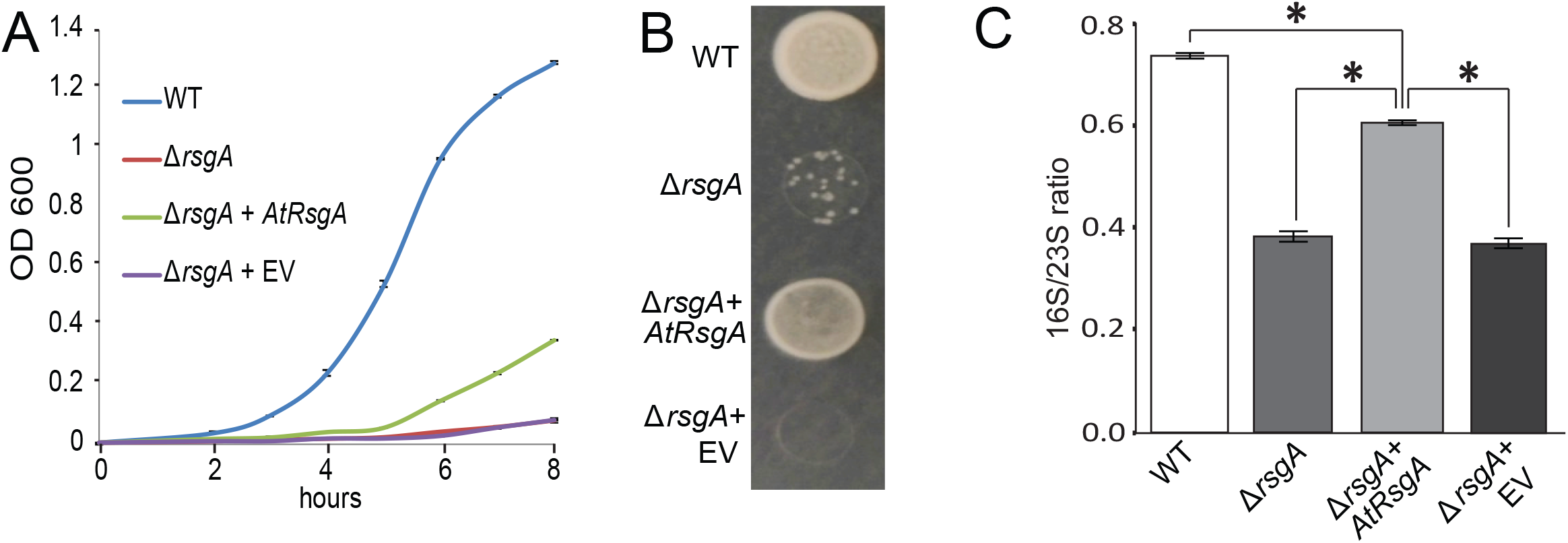
AtRsgA from *Arabidopsis thaliana* partially complements *ΔrsgA* deletion strain from *E.coli*. A) Growth curve of wild type (WT), mutant (*ΔrsgA*), complemented (*ΔrsgA+AtRsgA*) and control (*ΔrsgA+empty vector, EV*) strains in liquid media. The error bars indicate standard deviation (n = 3). B) Growth of transformed and non-transformed cultures on solid LB medium. C) 16S/23S rRNA ratios (as calculated from bioanalyzer electropherogram) of transformed and nontransformed cultures grown in liquid cultures. The asterisks indicate ratios significantly different from the complemented strain (*ΔrsgA+AtRsgA*, *P* < 0.05).

### Arabidopsis RsgA is essential for morphogenesis and viability

To decipher the potential role of *AtRsgA* in *Arabidopsis thaliana*, we analysed two transfer DNA (T-DNA) insertion lines, *rsgA-i* (insertion in the fourth intron) and *rsgA-e* (insertion in the third exon, Figure 2A). We could not obtain homozygous plants for the *rsgA-e* allele, and heterozygous plants produced wild-type (WT):heterozygous plants in 1:2 ratio (31:62, as defined by PCR), indicating that the *rsgA-e* allele is recessive and embryo or seedling lethal in homozygous state. Real-time PCR analysis showed significantly reduced *AtRsgA* transcript in both lines, with homozygous *rsgA-i* showing a stronger decrease (Figure 2B). While the heterozygous *rsgA-e* plants showed a wild-type like morphology, the homozygous *rsgA-i* plants were severely dwarfed and chlorotic at the rosette stage (Figure 2C) and as mature plants (Figure 2D). The dwarfed growth of *rsgA-i* is marked with both reduced growth rate (Figure S3A), rosette area (Figure S3B), and a significant reduction in seed yield (Table S1, Figure 2E). Interestingly, although heterozygous *rsgA-e* plants showed no net reduction in seed yield, 24.9±2.7 % of immature seeds were white (Figure 2D-E, Table S1), suggesting that the white seeds were homozygous for the *rsgA-e* allele and not viable.

**Figure 2.**
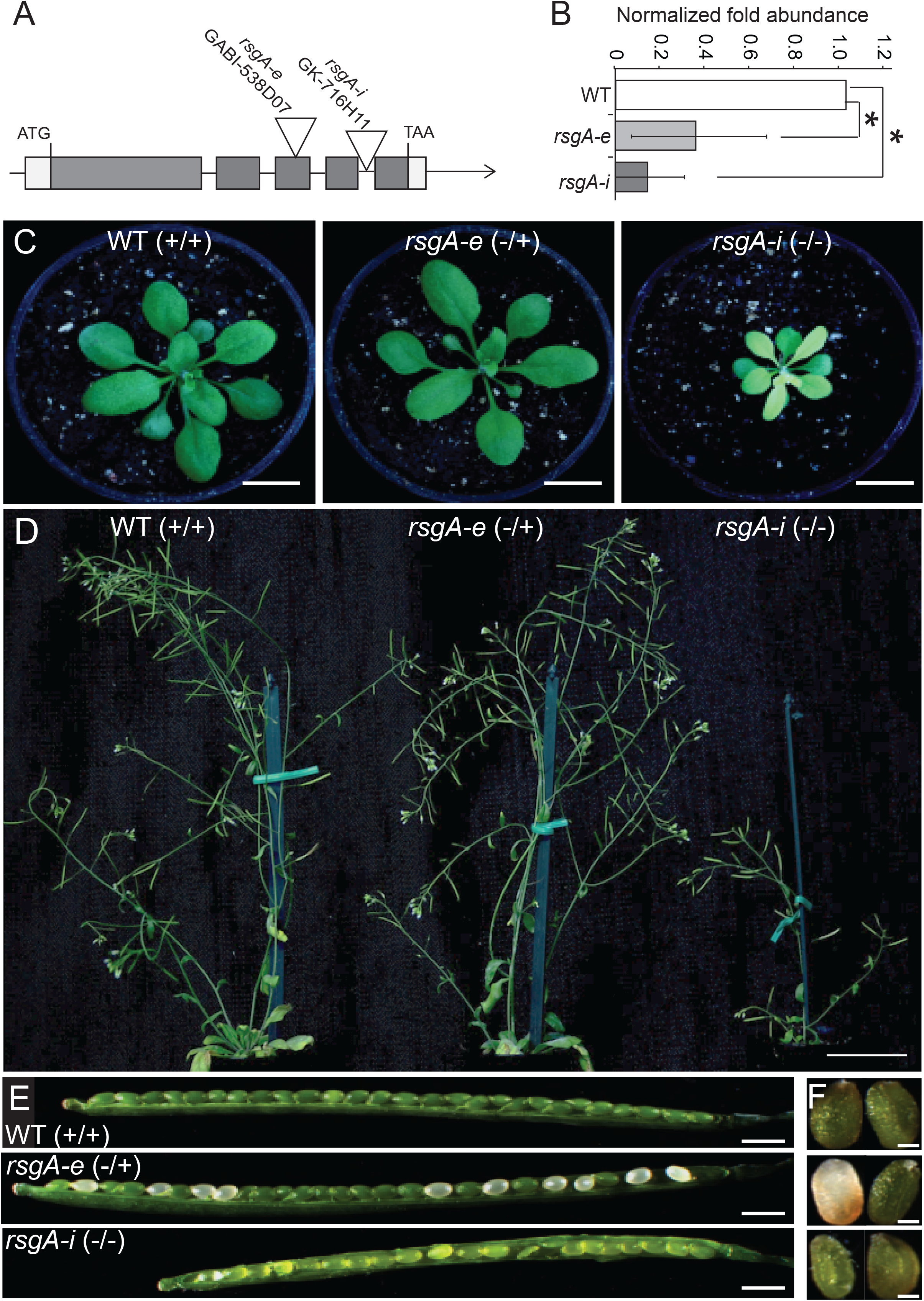
Phenotypes of *AtRsgA* T-DNA insertion lines in *Arabidopsis thaliana*. A) Exon-intron structure of *AtRsgA*, together with the position of the two T-DNA lines. Untranslated regions, exons and introns are indicated by white boxes, gray boxes and lines, respectively. B) *AtRsgA* expression analysis in heterozygous *rsgA-e* and homozygous *rsgA-i* lines. The expression of *AtRsgA* in *rsgA-e* and *rsgA-i* lines was normalized to its expression in WT sample. Error bars represent standard deviations (n=3, asterisk = P < 0.01). C) Rosette phenotype after four weeks of growth (scale bar = 1 cm). D) Phenotypes of eight week old plants (scale bar = 5 cm). E) Opened siliques from WT, *rsgA-e* and *rsgA-i* (scale bar = 1 mm). F) Exposed immature seeds from WT, *rsgA-e* and *rsgA-I* (scale bar = 500 μm).

To show that the T-DNA insertions in *AtRsgA* are causing the observed phenotypes, we analyzed progeny of heterozygous *rsgA-e* pollinated with homozygous *rsgA-i*, and observed the expected wild-type (WT/rsgA-i):dwarf (*rsgA-e/rsgA-i*) chlorotic ratio of phenotypes and genotypes (Table S2, Figure S4A, *X*^2^ P = 0.097 testing significantly different phenotype distribution). This showed that *AtRsgA* mutations are responsible for the observed phenotypes in both mutant alleles.

### Arabidopsis RsgA is involved in 30S ribosome maturation in chloroplasts

The chlorotic phenotype of *rsgA-i*, together with its ability to rescue a 30S assembly factor mutant in *E.coli* indicated that AtRsgA was involved in the maturation of the small subunit of chloroplasts. To verify this in plants, we compared rRNA profiles of *rsgA-i* and WT plants using a bioanalyzer electropherogram. The cytosolic plant rRNAs appear in form of two major peaks corresponding to 25S (60S subunit) and 18S (40S subunit) rRNA molecules, while the chloroplast rRNA profile is presented as a single peak for 16S rRNA (30S subunit) and three smaller peaks for 23S (23–1, 23–2, 23–3, 50S subunit) rRNA, which are generated post-transcriptionally by two hidden breaks (Figure S5, Figure 3A). The unchanged ratio of cytosolic subunits (25S/18S) in *rsgA-i* and WT samples indicated that the cytosolic ribosome biogenesis was not affected in *rsgA-i*. However, the 16S rRNA was significantly decreased as noted by decrease in 16S/23S-1 (Figure 3B, small and large chloroplast subunit) and 16S/18S (Figure 3C, small subunits from chloroplast and cytosol, respectively) amount in *rsgA-i* as compared to WT. The ratio of large chloroplast subunit was decreased when compared with the small cytosolic subunit (23S-1/18S, Figure 3C), but the change was less pronounced than for 16S.

**Figure 3.**
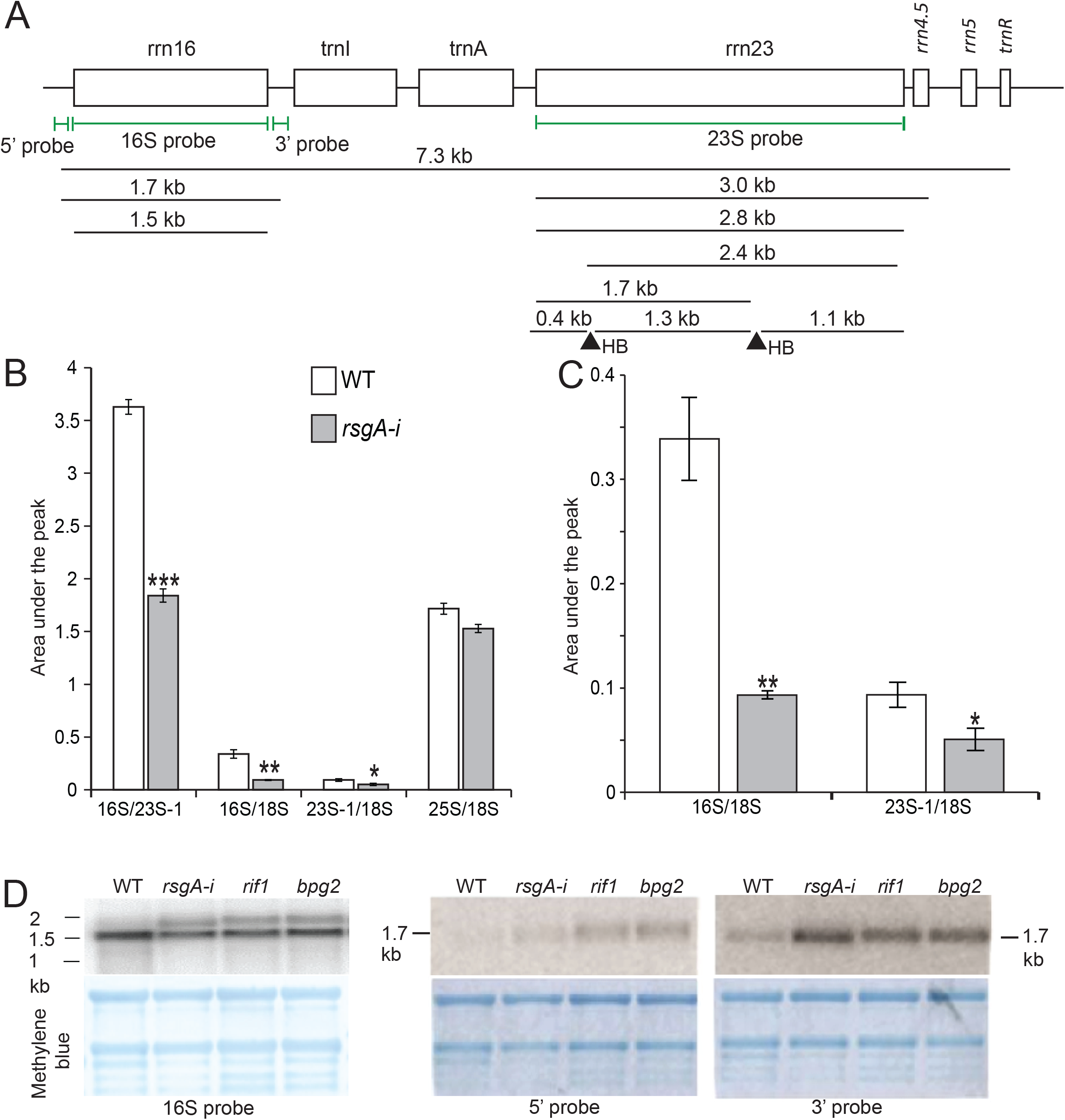
Northern blot analysis of *rsgA-i* and chloroplast 30S assembly mutants. A) Physical map of the plastid rDNA operon and maturation pattern of the plastid rRNA. The 7.3-kb primary transcript, processing intermediates, hidden breaks (HB, black triangles) and the mature forms of the 16S and 23S rRNAs are shown. Positions of hybridization probes are indicated by green lines. B) Bioanalyzer estimation of ratio of peak area of chloroplast rRNAs. Error bars indicate standard deviation (n = 3), while asterisk show significant difference between WT and *rsgA-i* (P < 0.01). C) Close-up of 16S/18S and 23S-1/18S ratios as estimated in (B). D) Northern blot analysis of mature 16S and 5’ and 3’ termini of 17S rRNA in WT, *rsgA-i, rif1* and *bpg2*. As a loading control, methylene blue-stained nylon membranes are shown. The bands on methylene blue membrane correspond to (from highest nt size): 25S, 18S rRNA from cytosolic ribosome, plastid 16S and two HB fragments of plastid 23S rRNA.

To corroborate the finding that 16S rRNA maturation was specifically affected and to identify the steps where the maturation pathway was perturbed, we performed Northern blot analysis using a specific probe against mature 16S rRNA (Figure 3A). As positive controls for small chloroplast subunit defects, we included two known chloroplast 30S assembly factor mutants *rif1* (resistant to inhibition by FSM)(Flores-Pérez *et al*., 2008) and *bpg2* (Brz-insensitive-pale green 2)(Kim *et al*., 2012). While in WT, the mature 16S rRNA was the most abundant rRNA, *rsgA-i, rif1* and *bpg2* mutants showed accumulation of 17S rRNA, a precursor of the 16S rRNA (Figure 3D)(Fristedt *et al*., 2014). Northern blot analysis showed normal processing for mitochondrial 18S (Figure S6A), mitochondrial 26S (Figure S6B), cytosolic 18S (Figure S6C) and chloroplast 23S rRNAs (Figure S6D), indicating that the *rsgA-i* defect is causing a specific maturation deficiency of chloroplast 16S rRNA. To ascertain if 17S rRNA precursor accumulation in *rsgA-i* was due to processing defect at 5’ and/or 3’ termini of 16S, we performed northern blot analysis using specific probes for the respective termini. The results showed that both 5’ and 3’ termini were not efficiently processed in the three mutants (Figure 3D).

Mutants of several *E.coli* ribosome assembly factors show temperature sensitivity (Charollais *et al*., 2003; Charollais *et al*., 2004; Connolly *et al*., 2008). Moreover, elevated temperatures can partially rescue lethality of chloroplast ribosome biogenesis defects in tobacco (Ehrnthaler *et al*., 2014). Indeed, *rsgA-i* mutants showed a more severe phenotype at lower temperature (18 °C) as compared to normal growth temperatures (23 °C, Figure S7). However, elevated temperature (28 °C) could not rescue the chlorotic phenotype observed at 23 °C (Figure S7A). Interestingly, although at lowered temperature (18 °C) *rsgA-i* seedlings exhibited more severe phenotype, this was not accompanied with visibly increased accumulation of 17S precursor, as compared to higher temperatures (23 °C and 28 °C, Figure S7B). Similar results were obtained for *rif1* and *bpg2* (Figure S7). Taken together, *rsgA-i, rif1* and *bpg2* are potentially active at a similar step of 30S subunit maturation.

To test if 17S rRNA can be integrated into mature ribosomes, we performed a polysome loading experiment. Polysomes consist of translating ribosomes in complex with mRNA, which can be separated from single ribosomes in a sucrose gradient upon centrifugation. Low sucrose fractions contain free subunits, monosomes and disomes, while high sucrose fractions contain polysomes, multiple ribosomes associated with one mRNA. While WT produced a single 16S rRNA in monosomes and polysomes (Figure 4A), the *rsgA-i* showed presence of 16S and 17S rRNA precursor in low (fractions 3-5) and high (fractions 11-12) density fractions (Figure 4B). The presence of 17S in high density fractions indicated that the immature 17S rRNA was incorporated into translating ribosomes. Furthermore, the lower ratio of 17S/16S in the high density fractions (Figure 4C), when compared to the low density fractions, indicated that ribosomes containing 17S rRNA tend to be less associated with mRNA.

**Figure 4.**
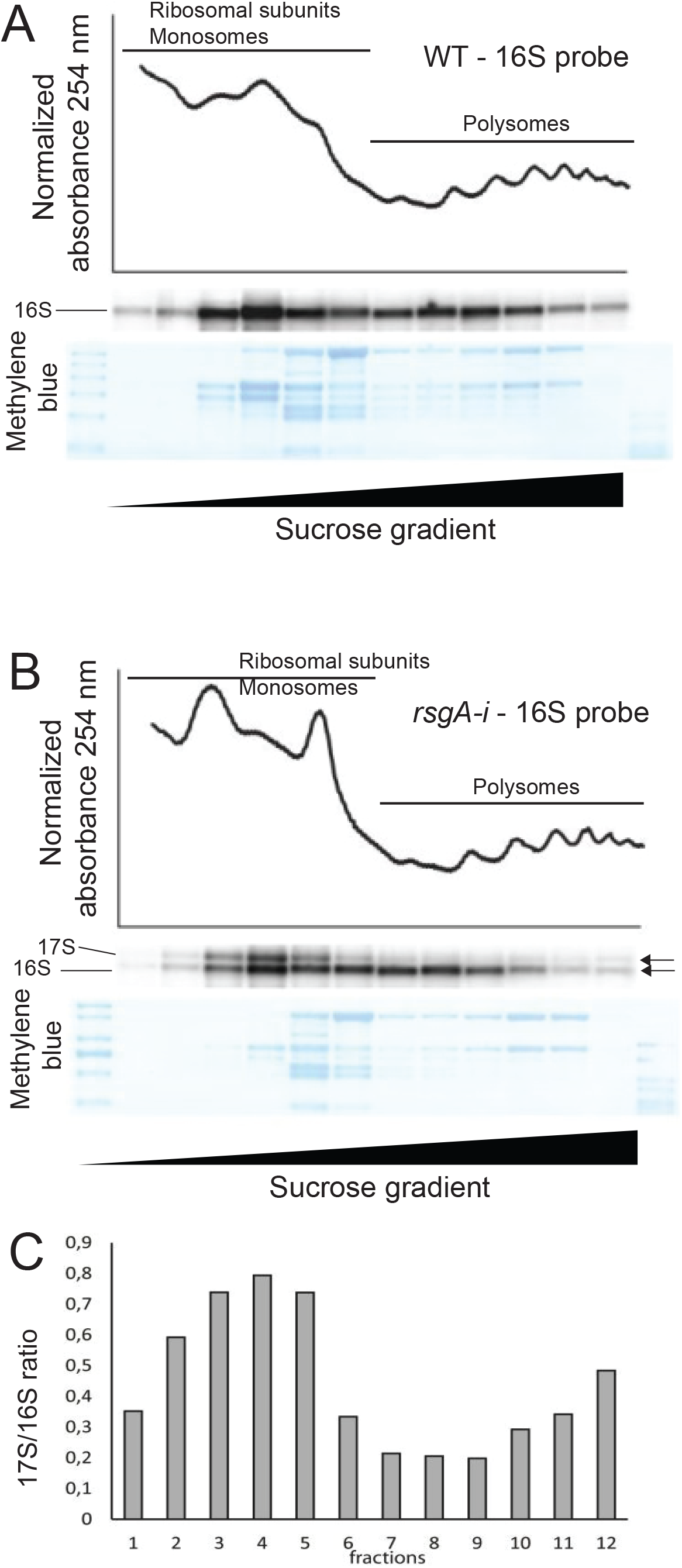
Polysome analysis of A) WT and B) *rsgA-i*. 254nm absorbance spectra were used to indicate the presence of ribosomal subunits, monosomes and polysomes. The collected fractions were subjected to 16S rRNA northern blot analysis, while methylene blue stain was used as loading control. The 17S and 16S rRNAs are indicated. C) Ratio of 17S/16S rRNA in the collected fractions.

### RsgA mutation leads to deficiency of the photosynthesis machinery

To corroborate the function of *AtRsgA* in chloroplast ribosome biogenesis, we performed *AtRsgA* expression analysis using Real-time PCR, and observed higher expression of the gene in green photosynthesizing organs as compared to non-photosynthesizing organs (roots and inflorescence), with additionally highest expression in mature seeds (Figure 5A). The expression of *AtRsgA* in green tissues was confirmed by high GUS (β-glucuronidase) activity in transgenics expressing GUS driven by the *AtRsgA* promoter (Figure 5B).

**Figure 5.**
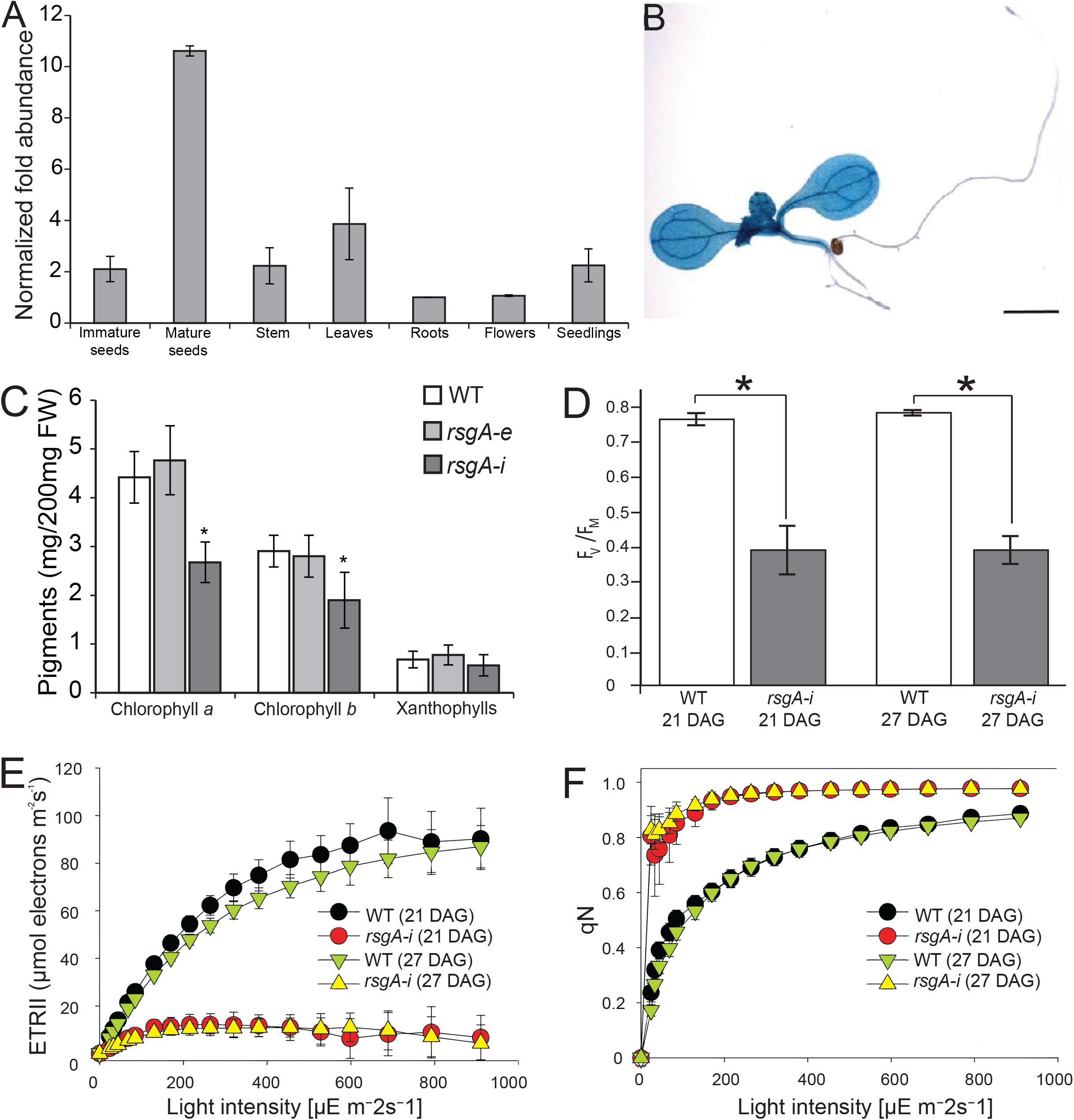
Expression analysis of *AtRsgA* and comparison of photosynthetic parameters of wild type and mutant lines. A) qRT-PCR of *AtRsgA* in different tissues, normalized to the expression in roots. B) *AtRsgA* promoter-GUS activity in one week old seedling showing high expression in photosynthetically active green tissue. Scale bar = 5mm. C) Photosynthetic pigment content measurements in 2-weeks old seedlings of WT, heterozygous *rsgA-e* and homozygous *rsgA-i* (n = 3, asterisk = P < 0.05). D-F) Photosynthesis efficiency parameters of WT and *rsgA-i* plants showing reduced photosynthetic capacity of *rsgA-i* plants. The measurements were taken at 21 and 27 days after germination (DAG). D) Maximum quantum efficiency of PSII. E) Light response curve of linear electron transport II (ETRII). F) Light response curve of non-photochemical quenching (qN).

To estimate the influence of the ribosome assembly defect on chloroplast function, we first estimated the levels of tetrapyrroles. Tetrapyrrole biosynthesis takes mainly place in chloroplast stroma, and through branching leads to production of chlorophyll a/b, hemes, siroheme, phytochromobilin and xanthophyll molecules (Mochizuki *et al*., 2010). Xanthophylls are most abundant in the light-harvesting complexes and their main function is photoprotection by channelling excess energy away from chlorophylls (reviewed in Ruiz-Sola and Rodríguez-Concepción, 2012). Interestingly, while the amount of chlorophylls was decreased in the *rsgA-i* mutant, xanthophylls remained unchanged, suggesting a differential regulation of the two compounds (Figure 5C).

Next, we measured several chlorophyll-a fluorescence parameters, such as rhe Fv/Fm ratio (maximum quantum efficiency of photosystem II photochemistry), linear electron transport rate (ETRII) and non-photochemical quenching (qN, dissipation of excess energy as heat). The results showed a decreased Fv/Fm ratio (Figure 5D) and linear electron transport capacity (Figure 5E), while the induction of non-photochemical quenching was strongly shifted to lower light intensities, and maximum non-photochemical quenching capacity was increased (Figure 5F), indicating reduced photosynthesis efficiency in *rsgA-i* plants.

### RsgA is important for proper plant and chloroplast morphology

To investigate the effect of potentially decreased chloroplast protein synthesis on chloroplast biogenesis, we investigated mesophyll cells of *rsgA-i* and WT seedlings by microscopy. The analysis revealed that chloroplasts in mesophyll cells of *rsgA-i* mutant plants were smaller than in WT (Figure 6A). We also observed that while the average cell size was not significantly different to WT plants in *rsgA-i*, chloroplast number per cell was significantly higher in *rsgA-i* (Figure 6B, P < 0.0001). Furthermore, despite the higher number per cell, the chloroplasts in *rsgA-i* line occupy less area in the cell due to their smaller size (Figure 6B).

**Figure 6.**
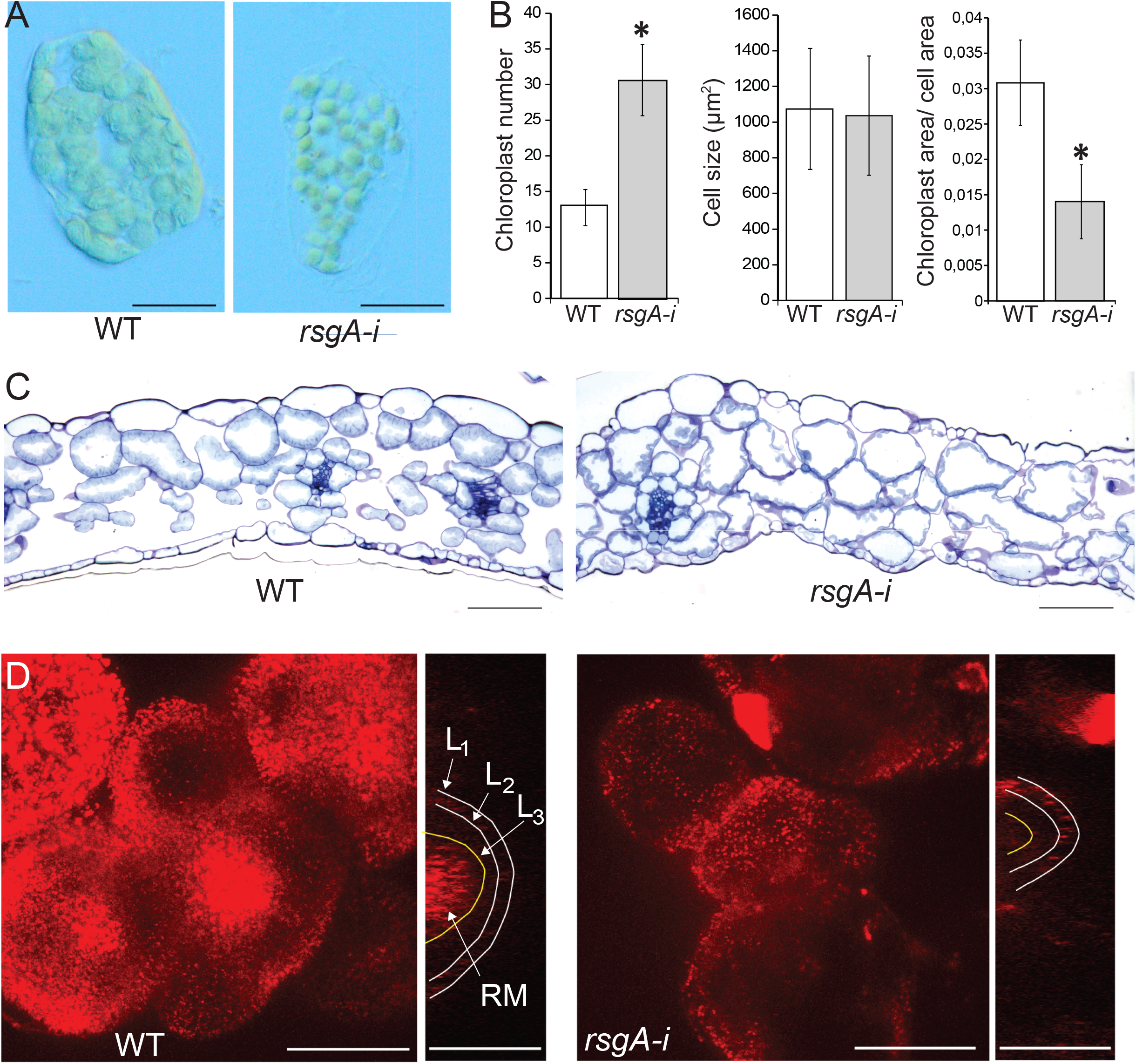
Physiological analysis of *rsgA-i*. A) Mesophyll cells from first true leaves of two weeks old WT and *rsgA-i* plants, fixed in 10 % formalin. Scale bar = 20 μm. B) Comparison of chloroplast number, mesophyll cell size and chloroplast area per cell area in WT (white bar) and *rsgA-i* (gray bar). Error bars represent standard deviation (n = 50, asterisk = P<0.01). C) Leaf cross-section with chloroplast stained with toluidine blue O (scale bar = 100 μm). D) Confocal image of shoot apical meristem and leaf primordia. Red color corresponds to plastid autofluorescence. The L1, L2 and L3 layers and rib meristem (RB) are indicated (scale bar = 50 μm).

Despite the similarity in mesophyll cell size, *rsgA-i* leaves were smaller compared to WT (Figure 2C, D), which could be caused by a decreased cell number and/or altered anatomy of the leaves. Therefore, we analyzed leaf cross-sections by microscopy, and in agreement with the previous result (Figure 6B), we observed no differences in cell sizes between *rsgA-i* and WT, and decreased chloroplast area occupancy in *rsgA-i* (Figure 6C). However, the mutant had visibly less intercellular air space between the mesophyll cells (Figure 6C), suggesting that increased cell packing might also play a role in the decreased leaf size.

In higher plants, most of the aboveground cells originate from a small group of undifferentiated, dividing cells called shoot apical meristem (SAM). The SAM consists of multiple cells layers (L_1-3_) and central rib meristem part. The shoot meristematic cells contain 10 to 20 proplastids (Possingham and Lawrence, 1983), which multiply to more than 100 chloroplasts in mesophyll cells of fully expanding leaves (Lopez-Juez and Pyke, 2005). To investigate if SAM morphology and chloroplast biogenesis were affected in *rsgA-i*, SAM of 4 weeks old plants was analyzed by confocal microscopy. We observed a severely reduced diameter of SAM in *rsgA-i* (Figure 6D). Furthermore, chlorophyll fluorescence, which is indicative of chloroplasts, was mostly located in rib meristem region in WT, while in *rsgA-i* the signal was largely absent (Figure 6D). Interestingly, proplastids accumulated more in some cells of L_1_ in *rsgA-i*, compared to WT plants (Figure 6D). These results indicate a connection between plastidial protein synthesis, chloroplast biogenesis and SAM formation.

### Impact of defective chloroplast protein synthesis on the transcriptome and proteome

A defect in plastid ribosome biogenesis machinery is expected to cause dramatic changes to the transcriptome and proteome of mutant plants, as it needs to adjust to the severely perturbed chloroplast function. To understand the molecular basis of this adjustment, we carried out transcriptomic (RNA sequencing) and proteomic (LC-MS/MS) analysis of *rsgA-i* and WT plants.

We first investigated how mRNA and protein expression of gene products targeted to different cellular compartments was perturbed in *rsgA-i*. The analysis revealed that most of the plastid-encoded transcripts tended to be up-regulated, while the corresponding proteins accumulated less (Figure 7A, Figure S8A for all compartments). This effect was also seen to a weaker degree for chloroplast-targeted nuclear-encoded genes. We also determined which gene products show significantly (P < 0.05) different expression in the mutant, and identified 1676 up-regulated and 1678 down-regulated transcripts, respectively. The proteomic analysis revealed 42 and 114 up- and down-regulated proteins, respectively (Table S3 for the transcriptome, and Table S4 for the proteome). Similarly to the previous result, the compartment with the largest number of differentially regulated elements was the chloroplast, where the majority of chloroplast-encoded and -targeted transcripts showed up-regulation. Conversely, the corresponding proteins showed significant decrease (Figure 7B, Figure S8B for all compartments). Indeed, this opposite behaviour of mRNAs and proteins corresponded to the largest fraction of differentially expressed proteins (Figure 7C, 64/156 proteins).

**Figure 7.**
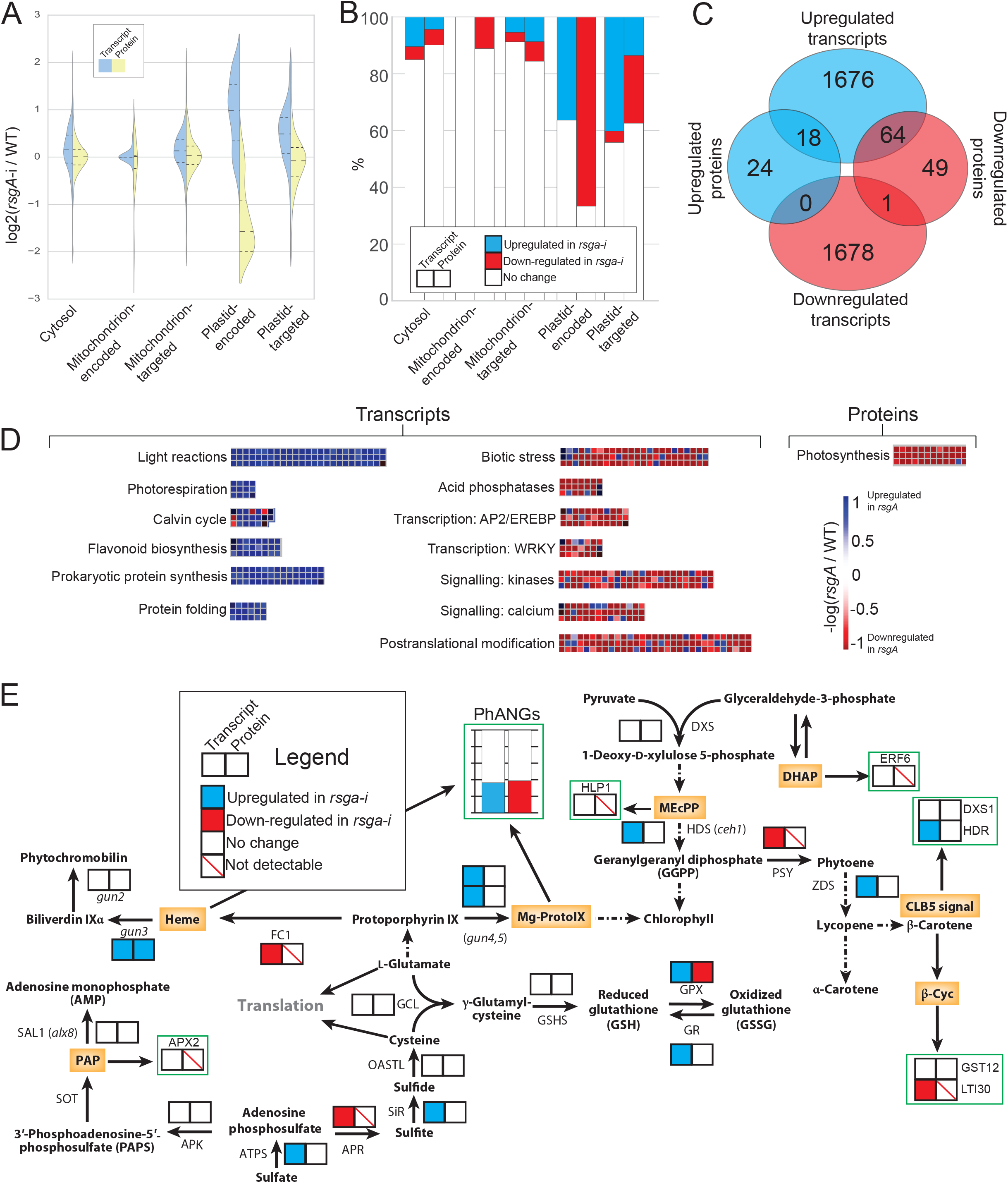
Transcriptome and proteome analysis of wild type and *rsgA-i*. A) Violin plots showing the log_2_-fold-change between *rsgA-i* and WT, for transcripts (blue side of the violin plot) and proteins (yellow side of the violin plot) targeted to, or encoded in the different compartments. The solid lines within the violin plot indicate median, while dashed lines show quartiles. Values above 0 indicate expression higher in *rsgA-i* than WT, while values lower than 0 indicate expression higher in WT than *rsgA-i*. B) Distribution of significantly (P < 0.05) differentially upregulated (blue bars), downregulated (red bars) or unchanged (white bars) transcripts (left bars) and proteins (right bars) in the different compartments. C) The numbers indicate how many proteins and transcripts are commonly up- or downregulated in *rsgA-i*, as compared to WT. D) MapMan analysis of the differentially expressed transcripts and proteins. Blue color indicates genes (cells in the heatmap) that are upregulated (blue) or downregulated (red) in *rsgA-i*. E) Biochemical pathways of the different metabolite retrograde signals. Known retrograde signals are shown in orange, solid arrows indicate a single biochemical step, and dash-dotted arrows are simplified representations of more than one biochemical step. Transcriptomic and proteomic responses are indicated by colored boxes, where blue, red, white and striked field indicates up-regulated, down-regulated, unchanged and not detectable, respectively.

To investigate which biological pathways were affected by the decreased protein synthesis in chloroplast, we performed a MapMan enrichment analysis (Thimm *et al*., 2004). Transcripts significantly upregulated in *rsgA-i* were involved in photosynthesis (light reaction, photorespiration, Calvin-Benson cycle) and chloroplast protein synthesis (Figure 7D, blue cells, P<0.05). Conversely, significantly downregulated transcripts belonged to pathways involved in biotic stresses, acid phosphatases, different transcription factor families, post-translational modification, and other signalling pathways (Figure 7D, red cells). MapMan analysis of the proteome revealed that photosynthesis was the only significantly enriched MapMan bin and most of these proteins were decreased (Figure 7D, P < 0.05). We confirmed these results by carrying out qRT-PCR and western blot analysis for some of the differentially expressed genes and proteins involved in photosynthesis. Indeed, transcripts of photosystem I (*psaA, psaB*), cytochrome b6 (*petA*), ATP synthase (*atpA*) were upregulated in *rsgA-i* (Figure S9A), while AtpA, PsaA and RbcL (RuBisCO) proteins were decreased (Figure S9B). Taken together, these results indicate that the decreased protein synthesis in chloroplast is met with an upregulation of photosynthesis- and protein synthesis-related transcripts.

### Analysis of chloroplast retrograde signalling pathways

To better understand the mechanism that induced higher expression of nuclear-encoded chloroplast-targeted transcripts and increased chloroplast number, we studied the retrograde signalling pathways. These pathways can control expression of nuclear genes by metabolic and protein signals to preserve plastid homeostasis in changing environment (operational signalling) and during development (biogenic signalling, reviewed in Chan *et al*., 2016). Still unknown signals from chloroplasts are conveyed to the nucleus, where they modulate the expression of photosynthesis-associated nuclear genes (PhANGs), plastid redox–associated nuclear genes (PRANGs), singlet oxygen–responsive genes (SORGs) and defense response genes. The proposed metabolic signals (orange boxes) and their biosynthesis pathways and genes they regulate (green boxes) are shown on Figure 7E.

Metabolic signals which are not likely candidates behind the observed phenotypes include 3-phosphoadenosine 5-phosphate (PAP), methylerythritol cyclodiphosphate (MEcPP), Heme and Dihydroxyacetone phosphate (DHAP). PAP alters gene expression by inhibiting the RNA-degrading activity of XRNs which target APX2 (Estavillo *et al*., 2011), whose transcript accumulation was unchanged in *rsgA-i* (Figure 7E). MEcPP induces wounding and high light stress-related nuclear gene expression of the chloroplast-targeted hydroxyperoxide lyase (HPL1) protein (Xiao *et al*., 2012), which was unchanged in *rsgA-i* (Figure 7E). Heme acts as a positive retrograde signal from plastids, and overexpression of Ferrochelatase 1 (FC1) transcript increases expression of PhANGs (Woodson *et al*., 2011). However, *rsgA-i* showed downregulation of FC1, while expression of PhANGS was increased (Figure 7E). DHAP activates MPK6, which subsequently induces the nuclear transcription of ERF-TFs (such as *ERF6*, Vogel *et al*., 2014), which remained unchanged in *rsgA-i*.

The metabolic signals that could cause the observed phenotypes in the *rsgA-i* mutant include Mg–protoporphyrin IX (Mg-ProtoIX), chloroplast biogenesis 5 (CLB5) and beta-cyclocitral (β-cyc). Mg-ProtoIX is an intermediate in chlorophyll biosynthesis (Kim *et al*., 2013) and might act by suppressing PhANG expression by binding to Long Hypocotyl 5 (HY) via Heat Shock 90 (HSP90)(Kindgren *et al*., 2011). Chlorophyll (and likely Mg-ProtoIX) is decreased in *rsgA-i*, while expression of multiple PhANGs is increased (Figure 7E). Loss of CLB5 causes a decrease in carotenoid biosynthesis, and repression of genes involved in the biosynthesis of isoprenoid precursors (DXS1 and HDR)(Avendano-Vazquez *et al*., 2014). The latter showed increased mRNA levels in *rsgA-i*. Finally, β-cyc expression increases at elevated ^1^O_2_ and induces expression of multiple genes, such as GST12 and LTI30 (Ramel *et al*., 2012), of which LTI30 was downregulated on the transcript level.

The protein signals are shown on Figure S10, together with expression analysis of 121 PhANGs (Zones *et al*., 2015), 418 SORGs (Gonzalez-Perez *et al*., 2011), 27 defense genes (Desveaux *et al*., 2004), 2 PRANGs (Chan *et al*., 2016) and 120 chloroplast-encoded genes (Table S3 for transcripts, Table S4 for proteins). The majority of the plastid-targeted nuclear-encoded genes were upregulated on the transcript level, while no changes were observed on the level of proteins (e.g. PRIN2, SG1, HDS). PhANGs and chloroplast-encoded genes were upregulated on the level of transcripts, but down-regulated on the level of proteins (39 and 59 proteins for PhANGs and chloroplast-encoded proteins analysed, respectively). SORGs and defense genes were down-regulated on the level of transcripts, with no observable differences in protein levels (397 and 17 proteins analysed for SORGs and defense genes, respectively). PRANG marker genes (APX2 and ERF6) showed no changes on the level of transcripts, indicating that the PRANG signalling pathway is not activated in *rsgA-i*. Of the nucleus-localized signals, only GLK transcription factors showed differential transcript expression. GLK1 and GLK2 induce expression of PhANGs and are down-regulated during chemically-inhibited plastid protein synthesis (Waters *et al*., 2009). Surprisingly, GLK2 was upregulated on the transcriptional level in *rsgA-i*, suggesting that GLK2 might be responsible for higher expression of PhANGs.

Several genetic perturbations can alter chloroplast numbers. These include RSH3, a guanosine penta- and tetraphosphate synthase, whose overexpression causes an increased number of chloroplasts, but decreased chlorophyll level and chloroplast diameter (Sugliani *et al*., 2016). Plastid division (PDV) mutants display lowered number of large chloroplasts, while the over-expression line shows a higher number of smaller chloroplasts (Okazaki *et al*., 2009). Finally, gibberellin (GA) deficient mutants have increased number of chloroplasts and chlorophyll, and similarly to *rsgA-i*, less air space between cells. However, the expression level of RSH3, PDV and GA-regulated genes, such as GNC and GNL (Figure S10)(Park *et al*., 2013), remained unaltered in *rsgA-i* suggesting that increased chloroplast number is potentially regulated by another, yet unknown mechanism.

## Discussion

This work elucidates the function of the Arabidopsis AtRsgA protein, as a novel chloroplast ribosome assembly factor. Despite more than 1.6 billion of years since the endosymbiotic event that gave rise to chloroplasts, AtRsgA could partially complement the slow growth and defect in 30S maturation of *E.coli ΔrsgA* mutant (Figure 1). The incomplete complementation could be attributed to the fact that AtRsgA was not codon-optimized for *E.coli*, due to minor structural differences between the *E.coli* and chloroplast ribosomes, or differences in pH, GTP availability and temperature between the cytosol of bacteria and plants. While *E.coli ΔrsgA* is viable (Figure 1)(Thurlow *et al*., 2016), the Arabidopsis *AtRsgA* is essential, as we could not obtain homozygous T-DNA insertion mutants of *rsgA-e* (Figure 2E). This suggests that the ribosome assembly (or the requirement of having a certain pool of functional ribosomes) is in some aspects different between *E.coli* and chloroplasts. Indeed, RbfA, an interaction partner of RsgA, shows processing defects in the 5’ end region of the 16S rRNA in *E.coli* (Xia *et al*., 2003; Datta *et al*., 2007), while the chloroplast counterpart shows defects in both 5’ and 3’ end processing (Fristedt *et al*., 2014). This suggests that ribosome assembly can be modified, and the assembly factors can gain or lose certain functions. The presence of a plant-specific region in the GTPase domain is especially interesting in this regard (Figure S1), and identification of its function warrants further investigation.

The *rsgA-i* intron mutant showed a typical chloroplast deficiency phenotype, such as chlorotic leaves with reduced rosette area (Figure 2), depleted photosynthetic pigments and reduced photosynthesis (Figure 5). These phenotypes have been described for numerous mutants with disturbed chloroplast biogenesis, chlorophyll production (Liu *et al*., 2015; Motohashi *et al*., 2007; Maekawa *et al*., 2015) and chloroplast ribosome assembly (Fristedt *et al*., 2014; Liu *et al*., 2010; Komatsu *et al*., 2010). These defects ultimately resulted in a massive growth retardation and decrease in rosette area of the mutant (Figure S3). Mechanistically, the *rsgA-i* mutant displayed a specific defect of the 16S rRNA accumulation that was accompanied by an over accumulation of 17S precursor in chloroplasts, whereas rRNA accumulation of cytosolic and mitochondrial ribosomes was unaffected. Furthermore, AtRsgA was shown to interact with RbfA, which is absent in mitochondria (Goto *et al*., 2011; Fristedt *et al*., 2014). Together, these evidences strongly suggest that AtRsgA is important for maturation of the small chloroplast subunit (Figure 3). While *rsgA-i* also shows a modest, but significant decrease in 23S rRNA (50S subunit, Figure 3C), we have demonstrated that no precursors accumulated in processing pathways of rRNAs from chloroplast 23S rRNA. This suggests that the free 50S subunit can be degraded either (i) due to a lower level of the 30S subunit or (ii) by decreased translation of chloroplast-encoded components of 50S (such as L2 and L22). Interestingly, *rbf1* which acts at a similar step of 30S subunit maturation (Figure 3D), showed no differences in 50S subunit accumulation (Fristedt *et al*., 2014), suggesting some dissimilarity in the role of these two proteins.

The negative correlation between chloroplast number and chloroplast size has been well known for over a decade (Figure 5 A,B), but the sensing and actuating mechanism underlying this phenomenon is largely unknown (Pyke, 1997). Similarly to *rsgA-i*, the RSH3 over-expression (controlling homeostasis of penta- and tetra-phosphate)(Sugliani *et al*., 2016) and PDV over-expression (Plastid Division Protein)(Okazaki *et al*., 2009) result in a larger number of smaller chloroplasts. However, transcripts of these genes did not show a significantly altered expression (Figure 7), suggesting that the herein observed increased chloroplast number is controlled by another mechanism.

Nuclear- and chloroplast-encoded photosynthetic proteins were significantly less abundant in *rsgA-i* (Figure 6C, D). While decrease of chloroplast-encoded proteins could be explained by the decreased protein synthesis capacity in chloroplast, the decrease of the nuclear-encoded photosynthesis-related proteins took place despite the increase of the corresponding transcripts. This degradation most likely takes place in chloroplasts rather than cytosol, and we suggest that it is caused by the inability to assemble chloroplast complexes that consist of both nuclear- and chloroplast-encoded subunits. Indeed, damaged and unassembled protein complexes have been shown to be rapidly degraded in chloroplasts (Adam and Ostersetzer, 2001).

Transcriptomic analysis of *rsgA-i* showed a potential compensatory response of protein translation deficiency in chloroplast, by upregulation of chloroplast- and photosynthesis-associated nuclear genes (PhANGs, Figure 7E, Figure S10). To better understand which signalling pathway induced the expression of photosynthesis-related transcripts in the nucleus, we analysed the metabolic and protein components of the retrograde signalling pathways. The analysis of differentially expressed genes revealed that the altered metabolic signal candidates are Mg–protoporphyrin IX (precursor of chlorophylls), CLB5 or β-cyc. Mg-protoporphyrin signal IX is a likely candidate, as the metabolite acts as a suppressant of PhANG expression (Kindgren *et al*., 2011), and the chlorophyll (and most likely Mg-protoporphyrin IX) levels are decreased in *rsgA-i* (Figure 5C), while PhANGs are upregulated (Figure 7E, Figure S10). The *clb5* mutant shows downregulaton of photosynthesis-related genes (Avendano-Vazquez *et al*., 2014), while we found that *CLB5* transcript and PhANGs are upregulated in *rsgA-i* (Figure S10). Treatment of *rsgA-i* with products and inhibitors specific to these signalling pathways (e.g. feeding the plants with Mg-protoporphyrin IX to suppress PhANG expression or treatment of *clb5* with fluridone to suppress its chlorotic phenotype)(Avendano-Vazquez *et al*., 2014), could reveal which of these signals plays a role in the observed phenotypes.

Taken together, our work has identified a novel assembly factor needed for maturation of 30S ribosomal subunit in chloroplasts, while the transcriptomic and proteomic analysis revealed the responses underpinning the adaptation to a decreased chloroplast function.

## Materials and methods

### Cloning of *E.coli* complementation vector and Arabidopsis GFP constructs

*AtRsgA* cDNA fragment was amplified by PCR from ABRC clone (S80886) using a high-fidelity polymerase (Phusion; New England Biolabs) and primers rsgA_fwd and rsgA_rv/rsgA_rv (Table S5). The product was inserted into pENTR4 vector using restriction based cloning (NcoI/NotI restriction sites) strategy and shuttled to pET60 (for *E.coli* complementation) or pUBC/pUBN destination vector (for GFP fusion constructs) using Gateway cloning strategy. *AtRsgA* promoter region was amplified from *Arabidopsis thaliana* Col-0 genomic DNA using primers promotor_fwd and promotor_rv (Table S5), digested with BamHI and NcoI and ligated with T4 ligase (Roche) into pCambia 1305 vector.

### *E.coli ΔrsgA* mutant complementation

*ΔrsgA* mutant was transformed with pET60 vector with either pT7::GST-AtRsgA-His or pT7::GST-HIS (empty vector without ccdB cassette) expression cassette via electroporation. Three colonies from transformed/ non-transformed *ΔrsgA* and WT *E.coli* were inoculated in 5mL LB medium and grown overnight at 28°C and 180 RPM shaking. 50 μl of overnight cultures were used to inoculate 5 ml fresh LB medium without antibiotics to achieve starting culture OD_600_ = 0.005. Cultures were grown for 8 h at 28°C with 180 rpm shaking, and OD_600_ was measured every hour. For plate growth experiment, 10 μl of each starting culture (OD_600_ = 0.005) was plated on LB agar plates without antibiotics and grown overnight at 28°C.

### Plant growth conditions and genotyping

Two independent T-DNA lines, *rsgA-e* (GABI_538D07) and *rsgA-i* (GK-716H11) were used for characterization of *RsgA (At1g67440)*. Mutant plants were genotyped with primers *rsgA-e* fwd and *rsgA-e* rv for GABI_538D07, primers *rsgA-i* fwd and *rsgA-i* rv for GK-716H11, and T-DNA left border primer GABI LB for GABI lines (Table S1), with PCR parameters as recommended for GABI lines. For growth on sucrose-containing medium, seeds were surface sterilized and plated on Murashige and Skoog agar medium containing 1 % Suc (Murashige and Skoog, 1962) and grown in growth chamber under 16-h day (140 μmol m^-2^ s^-1^) / 8-h night regime. For growth in soil, two weeks old seedlings were transplanted into soil and grown in the greenhouse at 16-h day (250 μmol m^-2^ s^-1^) / 8-h night regime at 20 /16 °C (day/night).

### rRNA analysis with Bioanalyzer

Total RNA from *E.coli* was isolated using TRizol reagent (Thermo Fisher Scientific). RNA from seedlings was harvested using Spectrum Plant Total RNA Kit (Sigma-Aldrich). rRNA ratios were analyzed with an Agilent 2100 Bioanalyzer using the Agilent RNA 6000 nano kit following manufacturer’s instructions. rRNA ratios were determined as described previously (Tiller *et al*., 2012). Bacterial electropherogram curves for 16S and 17S rRNA overlap, due to which peak area calculation using Bioanalyzer software could not be used. Therefore, peak area was calculated with normalmixEM function from the R package mixtools, which was applied to the sampled data in order to deconvolute the peak as a mixture of two Gaussian distributions. Pairwise Student t-test was used to calculate the significance of the differences in the ratio of the areas.

### Northern blotting

Northern blot analysis was essentially performed as previously described (Beick *et al*., 2008). In bried, total RNA was separated on formaldehyde-containing 1.2 % (w/v) agarose gels and blotted onto Hybond N membranes (GE Healthcare). Hybridization probes (Table S6) were generated by PCR amplification and purified by agarose gel electrophoresis, or commercially synthesized and then labeled with [α-32P]dCTP by random priming (Multiprime DNA labelling system; GE Healthcare) or with [γ-32P]dATP by 5’ end phosphate exchange (T4 PNK, New England Biolabs), respectively. Hybridizations were performed at 65 °C.

### Polysome loading assay

The analysis was performed as in (Barkan, 1993), with gradient concentrations of 15 %, 30 %, 45 % and 60 %. Samples were ultracentrifuged in 9 ml tubes for 4 hours at 37000 rpm in 4□C.

### Quantitative real-time PCR

cDNA used for qRT-PCR was obtained via Reverse transcriptase polymerase chain reaction using SuperScript™ IV Reverse Transcriptase (Invitrogen). qRT-PCR was carried out using the Maxima SYBR^®^ Green/ROX qPCR Master Mix (Thermo Scientific, UK). The expression levels of three technical replicates for each three biological replicates were measured on One Step Plus real time PCR system (Applied Biosystems, Life Technologies, Germany). Equipment setup, data collection and analysis were done using ABI SDS 2.3 software (Applied Biosystems, Life Technologies, Germany). The relative expression of target genes was normalized separately to housekeeping genes *Ubiquitin10* (*UBQ10*) and *Actin*. The samples which gave similar relative expression to both reference genes were considered for analysis. Data were analyzed by Livak method (Livak and Schmittgen, 2001).

### Protein Isolation, Gel Electrophoretic Separation and Immunodetection

100 mg of plant tissue was ground to fine powder and 150 μl of extraction buffer (40 mM Tris-HCl pH 7.0, 4 % SDS, 5 % glycerol, 2x Protein inhibitor cocktail) was added. Samples were vortexed for 5 min and spun down at 13000g for 30 min at 12°C. Supernatant containing total protein extract was collected and used for further experiments. The protein content was measured using BCA assay kit (Thermo Fisher Scientific). Total protein extract was separated by SDS-PAGE (6 % stacking gel, 12 % separation gel), and the proteins were subsequently transferred to polyvinylidene difluoride membranes (Immobilone; Millipore). Membranes were blocked with 5 % milk for 1 hour at room temperature with gentle shaking and then incubated with the primary antibody overnight at 4 °C. Blotted membranes were washed five times for 5 minutes using Tris-buffered saline with 1 % (v/v) Tween 20 (TTBS) buffer and incubated at room temperature with horseradish peroxidase-conjugated secondary antibody for 1 hour. The membranes were then washed five times for 5 minutes with TTBS buffer, and the immunoreactive proteins were visualized following 5 minutes incubation with detection reagents from the SuperSignal WestPico horseradish peroxidase detection kit (Thermo Scientific). Quantification of the immunoblots was done using the Fujifilm LAS-1000 software.

### Measuring plant growth kinetics and photosynthesis parameters

Total rosette leaf area was measured with the IMAG3D Maxi-Version of the Imaging PAM M-series, which was equipped with a 3D scanner based on a structured illumination approach (Heinz Walz GmbH, Effeltrich). Rosette leaf area was determined 21, 24, 27 and 30 days after sowing. To determine chlorophyll *a* fluorescence parameters on intact rosettes, the Maxi-version of the Imaging-PAM M-series was used. Light response curves between 0 and 1000 μE m^-2^s^-1^ actinic light intensity were measured with measuring times per intensity ranging between 150 seconds under light-limited and 60 seconds under light-saturated conditions. Plants were dark-adapted for 30 min prior to all measurements. Linear electron transport (ETRII) was corrected for leaf absorbance.

### Leaf cross-section analysis

For leaf cross section, two weeks old seedlings were fixed with 4 % paraformaldehyde and 0.2 % glutaraldehyde by vacuum infiltration and incubated at 4 °C overnight. The samples were washed in 0.1M phosphate buffer (pH 7.4), and dehydrated in an ethanol series (30, 50, 70, 80, 90 and 100 %) with incubation times of 1 hour in each solution. Samples were infiltrated with Technovit 7100 resin (Heraeus Kulzer) for 24 hours, followed by polymerization at room temperature (Beeckman and Viane, 2000). Microscopic analysis was performed on 5 μm cross-sections cut using a rotary microtome (RM 2265; Leica) and placed on poly-L-lysine-coated glass slides (Sigma). The slides were dried at 37 °C for 2 hours on a heating plate, and stained with toluidine blue O (0.05 %). Finally, the sections were examined with motorized epi-fluorescence microscope (Olympus BX 61) using cellP software (Olympus).

### Shoot apical meristem imaging and transient expression in protoplasts

Cut shoot apical meristems from *rsgA-i* line and wild type were transferred to a fresh 2 % agarose in a small round petri dish and used for imaging. 0.1 % Propidium iodide (PI) solution was used to stain cell walls. To image the meristem, a 40X water-dipping lens was used. The stack of images was obtained with a resolution of 0.5μm along the Z-axis. 561nm laser was used to excite PI and emission wavelengths between 605 to 630nm were recorded on a Leica SP8 confocal microscope. Arabidopsis protoplast were isolated and transformed with 10 μg of plasmid (Wu *et al*., 2009), and observed with Leica DM6000B/SP5 confocal laser scanning microscope (Leica Microsystems). The GFP was excited with 488 nm wavelength and emission of GFP was collected between 500 and 520 nm.

### Pigment content estimation

For chlorophyll extraction, leaf discs from 6 weeks old plants were cut and crushed in 80 % acetone solution followed by 12-16 hour incubation at room temperature. The readings were taken in triplicates at 470 nm, 645 nm and 662 nm using ELISA plate reader (Molecular devices, Versamax microplate reader). Chlorophyll a, chlorophyll b and xanthophyll + carotenoid amounts were calculated as in (Fan, 1997).

### RNA-Seq analysis

Samples were processed using LSTrAP (Proost *et al*., 2017), using default settings. All samples passed LSTrAP’s quality control. Log_2_ fold changes, p-values and corrected *p*-values (BH) for genes differentially expressed when comparing mutants with WT plants were calculated using DESeq2 (package version 1.10.1)(Love *et al*., 2014). Significantly differentially expressed genes (P < 0.05 after correction) were selected and used for a MapMan pathway analysis (MapMan v 3.6.0RC1)(Thimm *et al*., 2004). MapMan bins significantly enriched for genes (P < 0.05 after Benjamini and Hochberg (BH) correction) in this subset were determined and stored (Table S7).

### Proteomic analysis

50 μg of total protein extract was digest using trypsin. Peptides were re-suspended in 40 μl 5 % v/v acetonitrile, 2 % v/v trifluoroacetic acid. Measurements were performed on a Q Exactive HF coupled with an Easy nLC1000 HPLC (Thermo Scientific). Samples were loaded onto Acclaim PepMap RSLC reversed-phase column (75 μm inner diameter, 25 cm long Thermo Scientific) at flow 0.4 μl min^-1^ in 3 % v/v acetonitrile, 0.5 % v/v acetic acid, and eluted with a 3 % to 30 % v/v acetonitrile concentration gradient over 200 min, 30 % to 40 % in 20 min and from 40 % to 60 % in 20 min and followed by a washing step with 90 % v/v acetonitrile for 5 min, at 0.3 μl/min. Peptide ions were detected in a full scan from mass-to-charge ratio 200 to 2000. MS/MS scans were performed for the 15 peptides with the highest MS signal. Quantitative analysis of MS/MS measurements was performed with the Progenesis QI software (Nonlinear Dynamics). Proteins were identified from spectra using Mascot (Matrix Science). A t-test was performed for each peptide and the resulting p-values were corrected for multiple testing using FDR (BH). For the MapMan plots, only significantly changed P-values (corrected P < 0.05) were included in the plots (Table S8).

## Acknowledgments

We would like to thank Profs. Staffan Persson, Mark Stitt and Ralph Bock for the helpful discussions and feedback. We would like to thank the Max Planck Society for funding. The authors declare no conflict of interest.

## Supplemental figures

Figure S1. Multiple sequence alignment of RsgA protein sequence.

Figure S2. Bioanalyzer analysis of rRNA from *E.coli*.

Figure S3. Rosette growth dynamics.

Figure S4. Crosses of *rsgA-e* and *rsgA-i*.

Figure S5. rRNA Electropherogram of *rsgA-i* and wild type.

Figure S6. Northern blot against mitochondrial, cytosolic and chloroplast rRNAs.

Figure S7. Phenotypes of mutant and WT lines at different growth temperatures.

Figure S8. Differential expression analysis of *rsgA-i* and WT.

Figure S9. qRT-PCR and western blot analysis of the selected genes.

Figure S10. Chloroplast retrograde signalling responses.

## Supplemental tables

Table S1. Number and phenotypes of seeds in WT and the mutant lines.

Table S2. Phenotypic segregation of the *rsgA-e/rsgA-i* cross.

Table S3. Transcriptomic analysis of *rsgA-i* and WT.

Table S4. Proteomic analysis of *rsgA-i* and WT.

Table S5. Primer sequences.

Table S6. Probes used for Northern blot analyses.

Table S7. MapMan term enrichment analysis of the transcriptome.

Table S8. MapMan term enrichment analysis of the proteome.

